# Cancer vaccine attenuates carcinogen induced head and neck cancer with impaired early T cell response

**DOI:** 10.1101/2024.06.26.600828

**Authors:** Michihisa Kono, Masahiro Rokugo, John D. Quadarella, Shin Saito, Hiroki Komatsuda, Cong Fu, Sook-Bin Woo, Ann Marie Egloff, Ravindra Uppaluri

**Affiliations:** Department of Medical Oncology, Dana-Farber Cancer Institute, Boston, MA, USA; Department of Otolaryngology – Head and Neck Surgery, Asahikawa Medical University, Asahikawa, Japan; Division of Oral Medicine and Dentistry, Dana-Farber Cancer Institute and Brigham & Women’s Hospital, Boston, Massachusetts; Department of Surgery/Otolaryngology, Brigham and Women’s Hospital, Boston, MA, USA

**Keywords:** Head and neck cancer, 4NQO carcinogenesis, Genetically engineered mouse model, Antigen-specific T cell, Impaired T cell response

## Abstract

Effective T cell immunotherapy requires understanding antigen-specific T cell development during tumorigenesis and immune surveillance. Here, we aimed to examine the dynamics of antigen-specific T cells from tumor initiation through progression in a tobacco carcinogen mimetic, 4-nitroquinoline-1-oxide (4NQO)-induced head and neck carcinogenesis model utilizing genetically engineered K5^CreERT/+^/ROSA^OVA-GFP^/p53^fl/fl^ (KOG) mice. Our findings showed that early ovalbumin (OVA) expression via direct lingual tamoxifen (T) did not impact cancer development and survival, by comparing mice with tongue epithelium expressing OVA (KOG/T/OVA^+^) to those without OVA (KOG/T/OVA^-^) controlled by doxycycline. This equivalent tumor growth cannot be attributed to the loss of OVA expression. Intriguingly, although OVA-specific T cells were initially generated in tumor-draining lymph nodes (TDLN), they became undetectable 3 weeks after tamoxifen injection. Moreover, therapeutic anti-PD-1 was unable to restore OVA-specific T cells in TDLN and did not yield anti-tumor activity. Remarkably, OVA synthetic long peptide (SLP) vaccine induced OVA-specific T cells in KOG/T/OVA+ mice, and the combination of SLP vaccine and anti-PD-1 significantly reduced tongue tumor burden and prolonged survival. This study highlights the role of impaired endogenous antigen-specific T cell responses in immune resistance in head and neck cancer and the potential of cancer vaccines to improve outcomes.

## BACKGROUND

Head and neck squamous cell carcinoma (HNSCC) represents the seventh most common cancer worldwide, with 890,000 new cases diagnosed and 450,000 deaths each year (1). Patients with human papillomavirus (HPV)-unrelated HNSCC, mainly caused by carcinogen exposure such as tobacco, continue to suffer from recurrences or metastases, resulting in poor outcomes (1). These cancers develop from pre-malignant dysplastic lesions progressing to invasive cancer in the mucosal lining of the head and neck. Although not an immune desert malignancy, immune checkpoint inhibitors (ICIs) that have been approved as first-line therapy for unresectable recurrent or metastatic HNSCC show modest response rates (2). Thus, a better understanding of head and neck cancer specific immune evasion mechanisms, including those occurring in pre-malignancy is urgently needed.

Dissection of anti-tumor immune responses has revealed numerous mechanisms contributing to immune resistance as codified in the cancer immunoediting hypothesis (3). HNSCCs have been shown to exhibit several immune evasion mechanisms some of which include 1. down regulation of tumor cell stimulator of interferon genes (STING) expression (4), 2. genomic alteration of antigen processing machinery (5–7) and 3. a reduced adaptive T cell response possibly due to immunosuppressive cellular influence (8, 9) and/ or dendritic cell dysfunction (10). The cancer immunoediting hypothesis posits that tumor recognition and elimination occur at the earliest stages of tumorigenesis, but the initial dynamics between oral neoplastic cells and antigen-specific T cells leading to immune evasion are poorly understood. A major limitation has been the inability to model interactions that retain the carcinogenic origin of these cancers while simultaneously dissecting antigen-specific responses.

Herein, we established a model of 4-nitroquinoline-1-oxide (4NQO) carcinogen driven HNSCC by administering the carcinogen to genetically engineered mice with tongue epithelium specific Ovalbumin(OVA) expression and Tp53 gene deletion. Through this model, we aimed to investigate how antigen expression impacts the progression from oral premalignancy to invasive HNSCC, focusing on the dynamics of antigen-specific T cells dynamics.

## MATERIALS AND METHODS

### Sex as a biological variant

Our study examined male and female animals, and similar findings are reported for both sexes.

### Mouse models

Rosa^OVA-GFP(OG)^/Tp53^f/f^ mice have been described and were obtained from Dr. David Denardo (11) and K5^Cre-ERT/+^/ mice were obtained from the Jackson Laboratory. Mice were bred to obtain K5^Cre-ERT/+^/ Rosa^OVA-GFP(OG)^/Tp53^f/f^ (KOG) on the C57BL/6J background. The OG cassette was bred to homozygosity. All experiments used 6–10-week-old, male and female mice and experimental groups were matched in age and the number of males and females. TCR-transgenic OT-1 mice (C57BL/6-Tg (TcraTcrb)1100Mjb) were purchased from the Jackson Laboratory and bred in-house. All mice were maintained under pathogen-free conditions, and all animal experiments were approved by the institutional animal care and use committee (IACUC) at the Dana-Farber Cancer Institute.

### Administration of 4NQO, doxycycline and tamoxifen

4NQO (Sigma Aldrich) was dissolved in DMSO at a concentration of 50 mg/mL and then diluted in sterile water to a final concentration of 100µg/mL. KOG mice were exposed to 4NQO in their drinking water for 8 weeks and then switched to normal drinking water. (12). 4NQO drinking water was used in disposable cages and replaced weekly. In some experiments, doxycycline (625 mg/kg) was administered at the same time as the 4NQO drinking water. Fort he tamoxifen injections, we used the injection methodology described by Carper et al(13). A total of 30µl of 1mM 4-OHT (4-Hydroxytamoxifen, Selleck) in corn oil was injected into the epithelium of the tongue 3 times every 2 days on day 3, 5, 7 after 4NQO and doxycycline administration began. Following exposure to 4NQO, mice were visually checked for tumor development and weighed at least weekly.

### Tissue harvest

The KOG mice were sacrificed for humane endpoints decided by IACUC guidelines. Harvested lymph nodes (LNs) and spleen were processed into single cell suspensions using microscope slide glasses. Harvested tongues and tumors were dissociated using razor blades and then incubated with RPMI (Life Technologies) with collagenase Type IV (100µg/ml, Life Technologies) for 30 min at 37°C. Tongues were fixed in 10% formalin for 24 h. Following fixation, the long-axis of tongues were sectioned along the long axis and embedded in paraffin.

### Macroscopic tumor measurement

Tumor-bearing mice were photographed under anesthesia by retracting the tongue anteriorly and capturing images of the dorsum. ImageJ software was used to measure the area of the tongue and tumor, and the percentage of tumor area to tongue was calculated. The number of tumors per tongue were also counted.

### Microscopic evaluation

FFPE sections were stained with hematoxylin and eosin (H&E) and histopathological evaluation was performed by a board certified oral pathologist (S.B.W.). The evaluating pathologist was blinded to the treatment arms. The epithelial hyperplasia score was categorized into four levels (0–3). Meanwhile, the histopathological scoring of the most severe lesion was stratified into six levels: 0 for Normal, 1 for Atypia, 2 for Mild Dysplasia, 3 for Moderate Dysplasia, 4 for Severe Dysplasia or Severe Papillary Dysplasia, and 5 for Squamous Cell Carcinoma (SCC).

### Establishment of organoids

The methods for organoid culture and the medium formulation were adapted from Karakasheva et al.(14) with specific modifications for our study needs. Tongues from KOG mice were harvested, and then the epithelium was isolated using scissors and a dissecting microscope. Tongue epitheliums were incubated in HBSS (Thermo Fisher Scientific) with dispase (1:5, Corning), Fungizone (1:500, Thermo Fisher Scientific), gentamycin (1:10000, Thermo Fisher Scientific), and penicillin streptomycin (1:1000, Thermo Fisher Scientific) for 30min at 37°C in a thermomixer (700-800RPM). Tissues were removed from HBSS, incubated with 0.25% trypsin EDTA (Life Technologies) for 10 minutes, filtered through a strainer with soybean trypsin inhibitor (Sigma-Aldrich) and centrifuged. A similar procedure was used to make tumors organoids. Single cells (2000) were resuspended in 30µl Matrigel Matrix (Corning) with organoid media (1:1) and droplets were plated in a pre-warmed 24 well plate (Thermo Fisher Scientific). After incubation of the droplets at 37°C for 15min, pre-warmed organoid media was added. Organoid media was constituted of advanced DMEM/F12 (Thermo Fisher Scientific) with WRN conditioned media (3:100 Wnt/Noggin/R-Spondin), penicillin streptomycin (1:100, Thermo Fisher Scientific), HEPES (1:100, Life Technologies), Glutamax (1:100, Life Technologies), Gibco B-27 Supplement (2:100, Life Technologies), Gibco N-2 Supplement (1:100, Life Technologies), N-Acetylcysteine (0.5M, 2:1000, Fisher Scientific), mEGF (500µg/ml, 1:20000, Life Technologies)

### Flow cytometry

Isolated cells were stained with 1:500 Zombie Aqua (BioLegend) in PBS for 15 min at room temperature (RT). For SIINFEKL-tetramer staining, washed cells were incubated with 1:1000 PE-conjugated SIINFEKL tetramer (NIH tetramer core) in PBS with 0.5% FBS and 2 mM EDTA for 15 min at RT. For HAAHAEINEA-tetramer staining, cells were incubated with 1:100 PE-conjugated HAAHAEINEA tetramer (NIH tetramer core) in PBS with 0.5% FBS and 2 mM EDTA for 60 min at 37°C. Cells were then resuspended Fc block (BioLegend) for 10 min at 4°C and stained with 1:200 anti-mouse antibodies (Abs) in PBS with 0.5% FBS and 2 mM EDTA for 20 min at 4°C. For organoid assessment, organoids were pretreated with 2.5 mM 4-OHT for 96h and then with 2 µg/mL doxycycline for 48 h. For H2kb-SIINFEKL staining, cells were pretreated with 100 U/mL IFN-γ for 24h and stained with 1:1000 H2kb-SIINFEKL Ab (BioLegend,Clone:25-D1.16). GFP-expressing cell lines were used for GFP compensation. Data were acquired on Miltenyi MACSQuantX flow cytometer and analyzed using FlowJo v10.8.1 (BD) software.

### ELISA

T cells from KOG mouse TDLNs (5 × 10^5^) were co-cultured with SIINFEKL peptide (1 μg/mL, Thermo Fisher Scientific) in a 96-well plate (Thermo Fisher Scientific). For the *in vitro* OT-1 co-culture assay, tumor cells (5×10^4^) were co-cultured with T cells (5×10^5^) from OT-1 mouse splenocytes. After 24h, the supernatant was measured by IFN-γ ELISA (R&D) following manufacturer’s instructions.

### Adoptive OT-1 cell transfer

CD8+ T cells from OT-1 mouse splenocytes were purified with CD8a+ isolation kit, mouse (Miltenyi Biotec) and then labeled with Carboxyfluorescein Succinimidyl Ester (CFSE) using CFSE Cell Division Tracker Kit (BioLegend) according to manufacturer’s instructions. OT-1 CD8+ T cells (5 × 10^5^) were retro-orbitally injected into tumor-bearing KOG mice. LNs were harvested on day 3 after transfer and the CFSE-labeled CD8+ OT-1 T cells were assessed by flow cytometry. CFSE-positive cells were divided into CFSE-high (before or during proliferation) and -low cells (post-activation and proliferation) and their percentages were compared.

### Vaccination

For peptide vaccine treatment, OVA-derived synthetic long peptide (SLP, 50 µg, SMLVLLPDEVSGLEQISIINFEKLTEWTS, Peptide2.0) was injected into buccal mucosa of tumor-bearing KOG mice 3 times every 3 days along with 100 μg polyI:C (Invitrogen) as adjuvant.

### Cell lines and inoculation

The mouse oral squamous cell carcinoma MOC1 was generated as previously described (15) (16) (17). The mKate2-SIINFEKL plasmid was kindly provided by Dr. Clint T. Allen (National Institutes of Health) and SIINFEKL-expressing MOC1 (MOC1OVA) was engineered as previously described (18) (10). These cell lines were tested for mycoplasma every 6 months. All cell lines were cultured with the 2:1 mixture of IMDM (Life technologies) and Ham’s-F12 nutrient mixture (Thermo Fisher Scientific) with 100 U/mL penicillin streptomycin (Thermo Fisher Scientific), 5% heat inactivated Fetal Bovine Serum (FBS) (Sigma Aldrich), 5 ng/mL Epidermal Growth Factor (EGF) (Life Technologies), 400 ng/mL hydrocortisone (Sigma Aldrich), and 5 µg/ml insulin (Sigma Aldrich). 3×10^6^ MOC1 cells were inoculated into the buccal mucosa of KOG mice or wildtype C57BL/6 mice.

### In vivo antibody treatment

Anti-PD-1 mAb (250 μg, clone: RMP1-14, BioXCell) or isotype control (BioXCell) was injected intraperitoneally into KOG mice. Anti-CD40L Ab (BioXCell, 250µg) was injected intraperitoneally on day 14.

### Statistical analysis

Statistical analyses were carried out using GraphPad Prism 9 software (GraphPad). Comparisons of two independent groups were performed using Student’s *t*-test and multiple comparisons were evaluated using ANOVA with Tukey’s multiple comparison adjustment. Kaplan–Meier curves and log rank tests were used to assess survival differences. Differences were considered statistically significant at two-sided p <0.05, marked as *p < .05, **p < .01, ***p < .001, and ns = not significant. All data are shown as means ±SEM.

## RESULTS

### Establishment of a HNSCC mouse model with tongue epithelium-specific OVA expression and p53 deletion

The development of a chemically induced HNSCC model using 4NQO, a carcinogen that induces tobacco-associated gene mutation signatures, poses significant challenges for tracking antigen-specific T cells due to tumor heterogeneity. To address these challenges, we developed a 4NQO-induced oral cancer model using genetically engineered K5^Cre-ERT/+^; Rosa^OVA-GFP(OG)^; p53^f/f^ (KOG) mice. The OG mice, previously utilized in pancreatic and lung cancer models (11) were engineered to enable tongue epithelial basal keratinocyte keratin 5 (K5)-specific expression of OVA antigen and knockout of p53 upon tamoxifen injection (Figure 1A). Antigen expression is turned off with doxycycline administration. By administering 4NQO to these mice, we established an HNSCC mouse model that closely mimics human HNSCC with specific antigen expression, as shown in Figure 1A.

**Figure 1:**
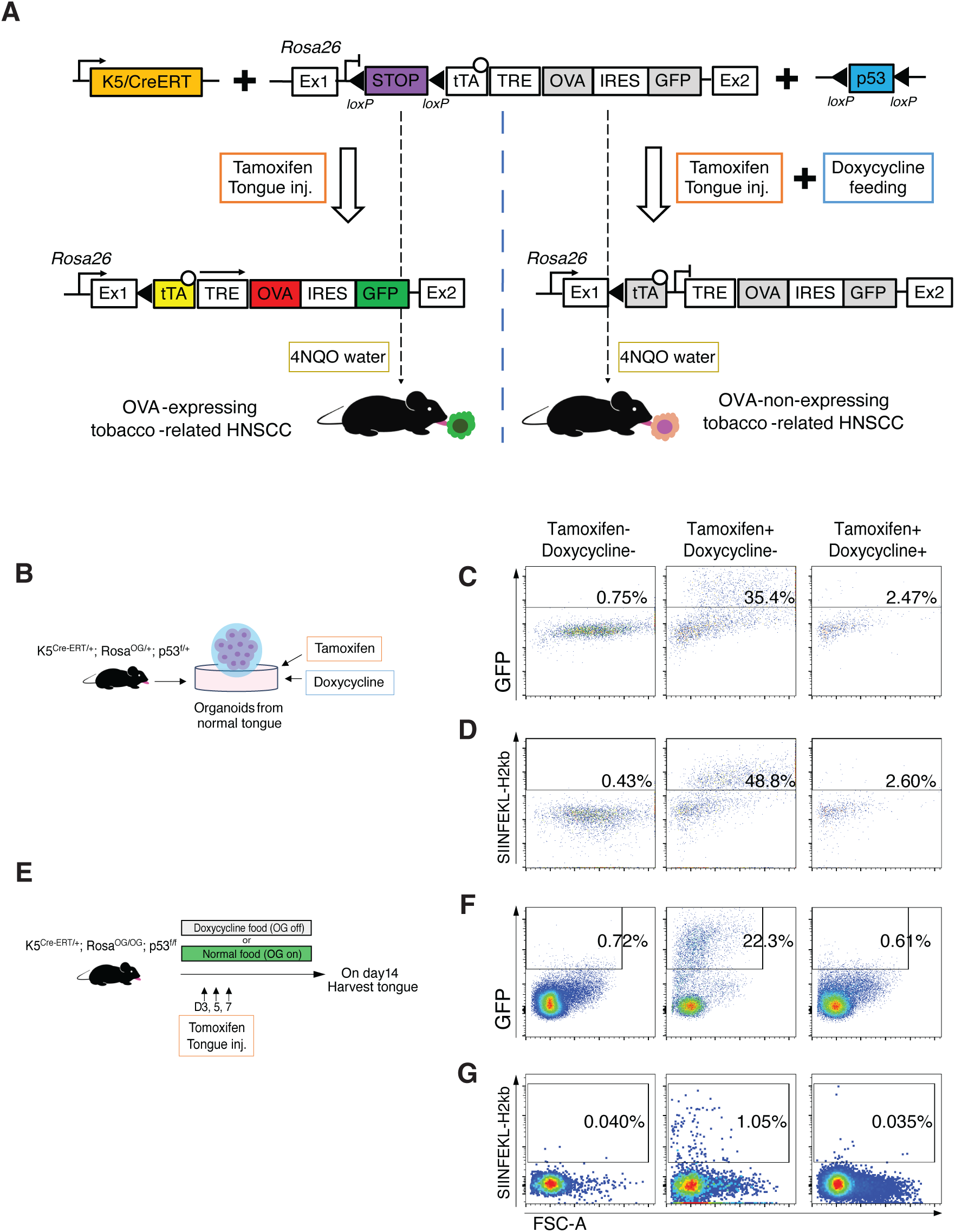
Establishment of KOG model. A. Genetic loci and strategy for KOG model. B. Experimental schema for C and D. Organoids were created from K5^Cre-ERT/+^; Rosa^OVA-GFP(OG)/+^; p53^f/+^ normal tongue and co-cultured with 4-hydroxytamoxifen (4-OHT, 2.5mM 96h) and then with doxycycline (2µg/ml 48h). For H2kb-SIINFEKL staining, cells were pretreated with 100U/mL IFN-γ for 24h. C and D. GFP expression (C) and SIINFEKL-H2kb expression (D) in KOG normal tongue organoids treated with 4-OHT and doxycycline. Data were pre-gated on live and single cells. Representative of 3 independent experiments. E. Experimental schema for F and G. K5^Cre-ERT/+^; Rosa^OG/OG^; p53^f/f^ mice were fed doxycycline or normal food and 4-OHT was injected into the tongue epitheliums 3 times every 2 days and harvested on day14. F and G. GFP expression (F) and SIINFEKL-H2kb expression (G) in KOG mouse tongue treated with 4-OHT and doxycycline. Data were pre-gated on live and single cells. Representative of 3 independent experiments.

To validate this genetic strategy, we first created organoids from the normal epithelium of K5^Cre-ERT/+^; Rosa^OG/+^; p53^f/+^ mice (Figure 1B). Organoids derived from normal tongue epithelium increased GFP and H2-K^b^-SIINFEKL expression upon tamoxifen treatment, and expression was abolished with doxycycline treatment (Figure 1C and D). Next, to confirm that this genetic modification works in vivo, tamoxifen was injected into the KOG mouse tongues 3 times every 2 days in the presence or absence of doxycycline. One week later, the tongues were harvested and single cells were analyzed by flow cytometry (Figure 1E). Results confirmed that GFP and H2-K^b^-SIINFEKL were also expressed by tamoxifen injection in vivo and eliminated by doxycycline feeding (Figure 1F and G). Together, we established a novel genetically engineered mouse with regulatable expression of tongue epithelium-specific OVA antigen .

### Early expression of OVA antigen did not affect 4NQO-induced HNSCC progression

Using the KOG mouse, we investigated how early antigen load affects carcinogen-induced tumorigenesis and anti-tumor immunity (Figure 2A). Mice were treated with 4NQO water for 8 weeks and then switched to normal water (Figure 2A). Lingual tamoxifen injection (T) to induce OVA expression and delete Tp53 was performed 3 days after the start of 4NQO water treatment (KOG/T/OVA^+^). A matched cohort was concurrently fed a doxycycline diet to turn off OVA expression (KOG/T/OVA^-^). Surprisingly, the tongues of KOG/T/OVA^+^ formed atypia and mild dysplasia at 9-15 weeks, and progressed to severe dysplasia and carcinoma at 23-30 weeks, comparable to KOG/T/OVA^-^ mice without elimination of antigen response (Figure 2B-D). At 22 weeks, macroscopic tumor area and number of tumors per mouse were determined to be comparable in both groups (Figure 2E, F, and S1). Similarly, overall survival rates and mouse weights showed no significant differences (Figures 2G and H). These results indicated that early expression of OVA antigen during tumorigenesis did not significantly affect tumor development in this model.

**Figure 2:**
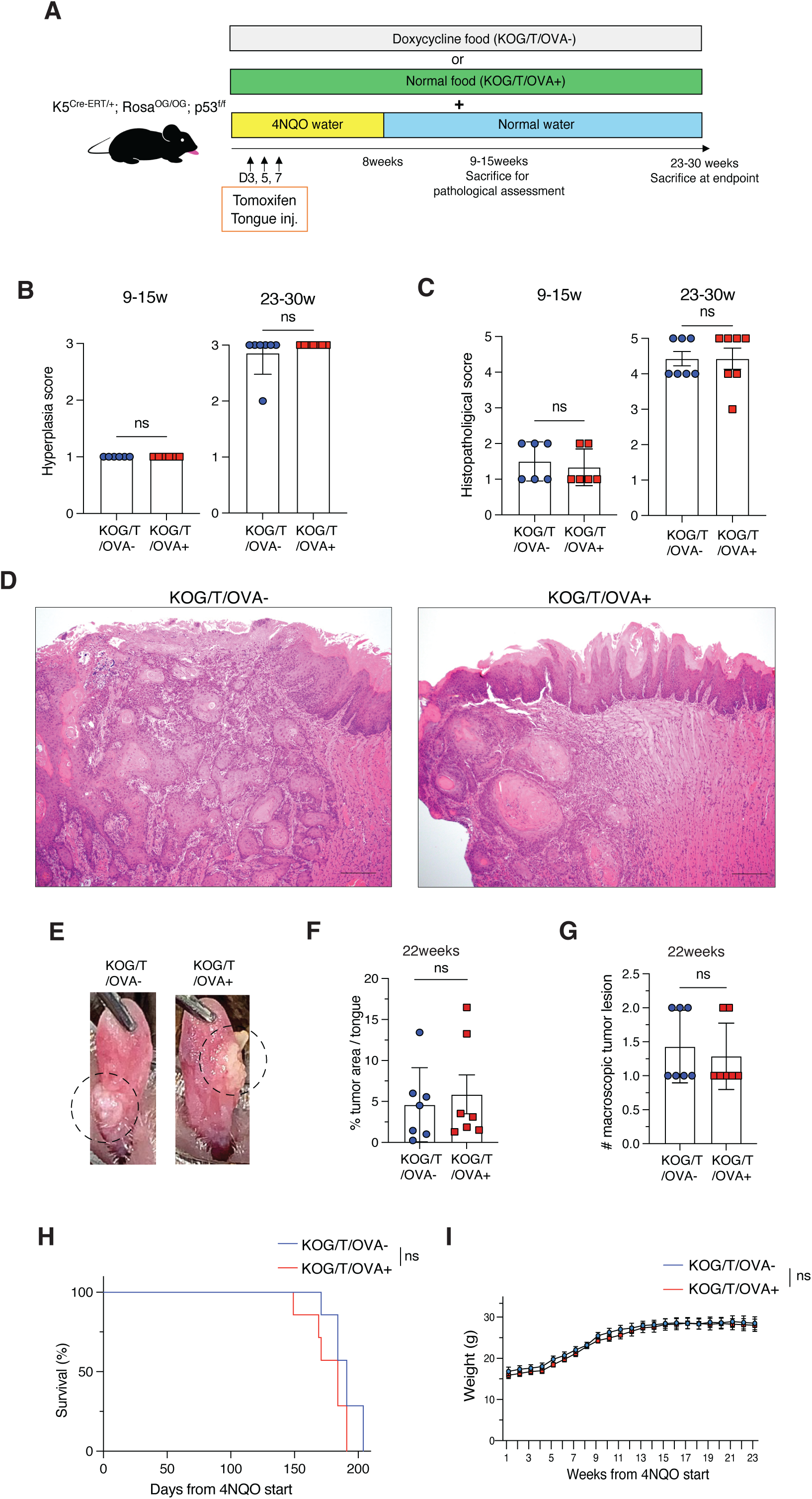
Early expression of OVA antigen did not affect 4NQO-induced HNSCC progression. A. Experimental schema for B-H. KOG mice were provided with 4NQO water for 8 weeks and then switched to normal water. Doxycycline or normal food was initiated concurrently with 4NQO. 4-OHT was injected on days 3, 5, and 7. Mice were sacrificed at specific time points or at the study endpoint. B. Epithelial hyperplasia scores on histopathological evaluation in KOG/T/OVA^+^ and KOG/T/OVA^-^ mice at 9-15 weeks and 23-30 weeks (means ± SEM). The epithelial hyperplasia score was categorized into four levels (0–3). C. Histopathological scores in KOG/T/OVA^+^ and KOG/T/OVA^-^ at 9-15 weeks and 23-30 weeks (means ± SEM). The histopathological scoring of the most severe lesion was categorized into six levels: 0 for Normal, 1 for Atypia, 2 for Mild Dysplasia, 3 for Moderate Dysplasia, 4 for Severe Dysplasia or Severe Papillary Dysplasia, and 5 for Squamous Cell Carcinoma (SCC). D. Representative SCC lesion of tumor-bearing tongues of KOG/T/OVA^+^ and KOG/T/OVA^-^ mice at 23-30 weeks. E. Representative photographs of tumor-bearing tongues of KOG/T/OVA^+^ and KOG/T/OVA^-^ mice at 23-30 weeks. F. Percentage of the macroscopic tumor area at 22 weeks (means ± SEM). G. The number of the macroscopic tumor lesion at 22 weeks (means ± SEM). H. Kaplan-Meier survival curve. I. Weekly body weight. *p<0.05, **p<0.01, ***p<0.001. Significance was evaluated by unpaired Student’s t-test. Kaplan– Meier curves were used for survival.

### OVA antigen is maintained on tumor cells during tumorigenesis

To determine whether the loss of OVA expression was responsible for the equivalent tumor growth in the comparative experiment, we first assessed GFP expression in tongue epithelium of KOG/T/OVA^+^ at 12-weeks and observed no loss of GFP fluorescence (Figure 3A). Second, organoids were created from 25-week-old KOG/T/OVA^+^ mice and we found that these cells continued to express GFP and also H2-K^b^-SIINFEKL (Figure 3B, C, and S2A). Third, confirming that tumor cells were recognized by OVA-specific T cells, OT-1 CD8+ T cell co-culture supernatants were positive for IFN-γ, indicating antigen specific OT-1 CD8 T cell activation (Figure 3D). Finally, we then examined whether antigen was processed and presented in tumor-draining lymph nodes (TDLNs). CFSE-labeled OT-1 CD8^+^ T cells were transferred into tumor-bearing KOG/T/OVA^+^ mice (20 weeks post-4NQO) and TDLNs were harvested 3 days after transfer (Figure 3E). OT-1 CD8+ T cells responded to antigen and proliferated in TDLNs (Figure 3F, G, and S2B). From these results, we concluded that OVA expression was maintained during 4NQO-induced tumor development in mice without doxycycline.

**Figure 3:**
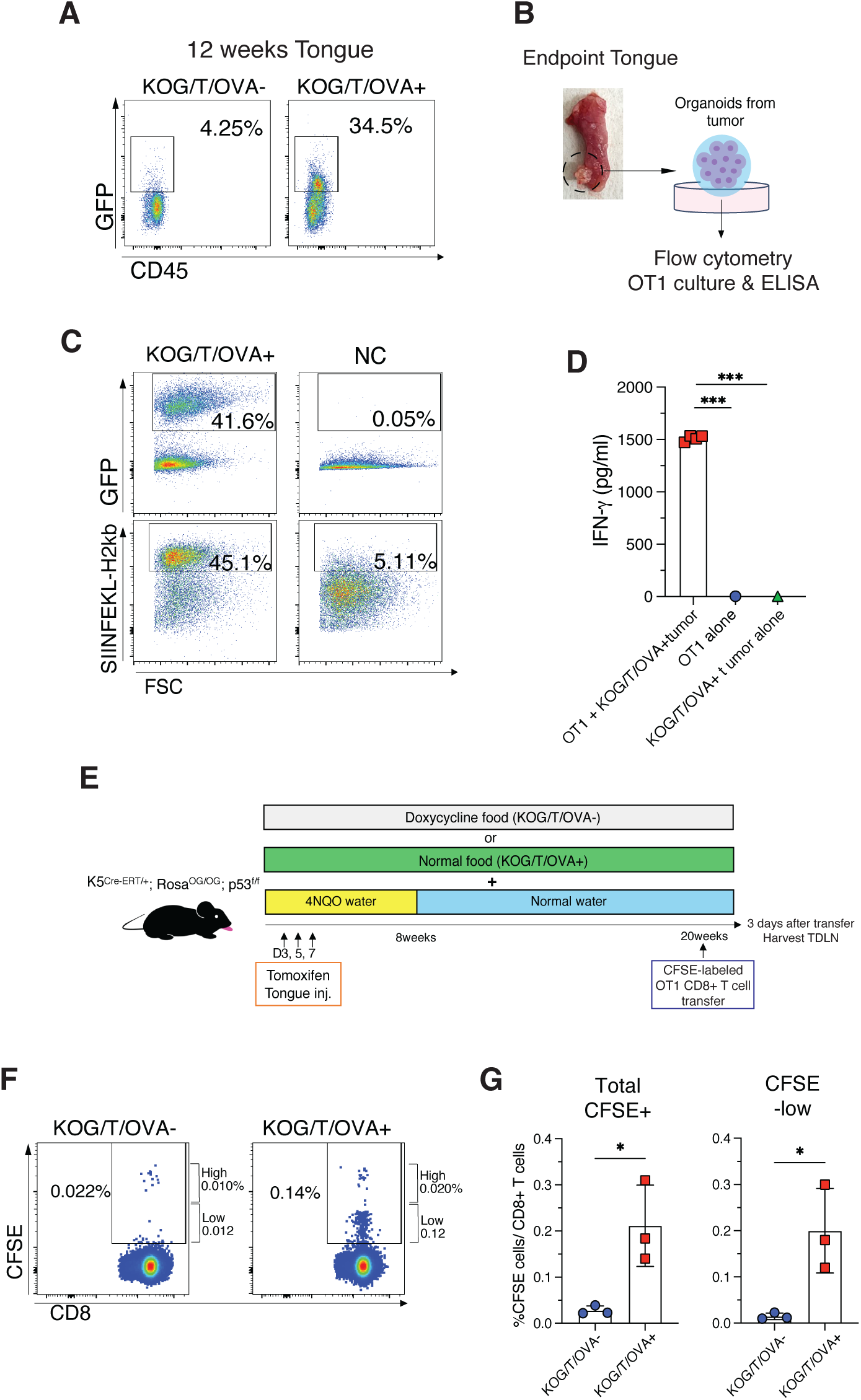
OVA expression was maintained on tumor cells during tumorigenesis. A. GFP expression of the tongue epithelium cells from 12-weeks KOG/T/OVA^+^ and KOG/T/OVA^-^mice. Data were pre-gated on live cells, single cells, and CD45-. Representative of three independent experiments. B. Experimental schema for C and D. Organoids were created from KOG/T/OVA^+^ tongue tumors and evaluated by flow cytometry and the experiment of OT-1 co-culture and ELISA experiments. C. GFP and SIINFEKL-H2kb expression of the tumor organoids from tumor-bearing mouse tongue. For assessment of SIINFEKL-H2kb expression, organoids were pre-treated with 100U IFN-γ for 48h. Data were pre-gated on live and single cells. Representative of 3 independent experiments. D. Quantification of IFN-γ production in supernatants of the cultures with tumor cells and OT-1 T cells (means ± SEM). Tumor organoids were dissociated into single cells and co-cultured with OT-1 T cells for 24 hours. The supernatants were assessed by IFN-γ ELISA. E. Experimental schema for F and G. CD8 T cells were purified from splenocytes of OT-1 mice and labeled with CFSE. CFSE-labeled OT-1 CD8+ T cells were transferred into tumor-bearing KOG mouse at 20 weeks and TDLNs were harvested 3 days after transfer. F. CFSE expression in tumor draining lymph nodes (TDLNs) 3 days after OT-1 transfer. Data were pre-gated on live cells, single cells, CD45+, CD3+, and CD8+ cells. CFSE-positive cells were divided into CFSE-high (before or during differentiation) and -low cells (post-differentiation). Representative of 2 independent experiments (n=3). G. Quantification of the percentage of total of CFSE-positive, CFSE-high, and CFSE-low cells in CD8+ T cells in TDLNs. Representative of 2 independent experiments (means ± SEM). *p<0.05, **p<0.01, ***p<0.001. Significance was evaluated by unpaired Student’s t-test.

### OVA-specific T cells were generated but subsequently became undetectable

Given that OVA antigen expression was maintained in the tongue epithelium and in tumor tissue during tumor progression, we hypothesized that the underlying cause of the lack of tumor regression by anti-tumor immunity was dysfunction within the T cell compartment. Thus, we chronologically assessed OVA-specific T cells in TDLNs throughout the 4NQO carcinogenesis protocol (Figure 4A). SIINFEKL-tetramer+ T cells were initially generated in TDLNs one week after the last injection of tamoxifen but surprisingly became undetectable three weeks later and remained absent through the remainder of observation and endpoint of study (Figure 4B, C, and S3A). Supernatants of TDLN cultures with SIINFEKL peptide were assayed by IFN-γ ELISA, and OVA-reactive T cells were initially detected ex vivo but no OVA-reactive T cells were detected by the 4-week timepoint (Figure 4D). OVA-specific CD4 T cells, which are not regulatory T cells, were also generated early on but were not detected later (Figure S3B). OVA-specific T cells remained lost even after the stimulation with a CD40 agonist, which was administered when these T cells were still present. (Figure S3C). To determine whether KOG mice have immune tolerance to OVA, we transplanted an OVA-expressing syngeneic mouse oral carcinoma (MOC1-OVA) into the buccal mucosa of KOG or wildtype C57BL/6 mice (Figure 4E). OVA-specific T cells were generated in KOG mouse tumors and TDLNs but their numbers were reduced compared to wildtype C57BL/6 mice (Figure 4F and G). These results suggest that KOG mice retain the cellular capacity to react to OVA antigen albeit at a reduced level compared to wild type mice and that tamoxifen-induced OVA-specific T cells became subsequently undetectable. Collectively, these data are consistent with impaired OVA-specific T cell response as the reason why antigen activated T cell response did not reduce tumor growth.

**Figure 4:**
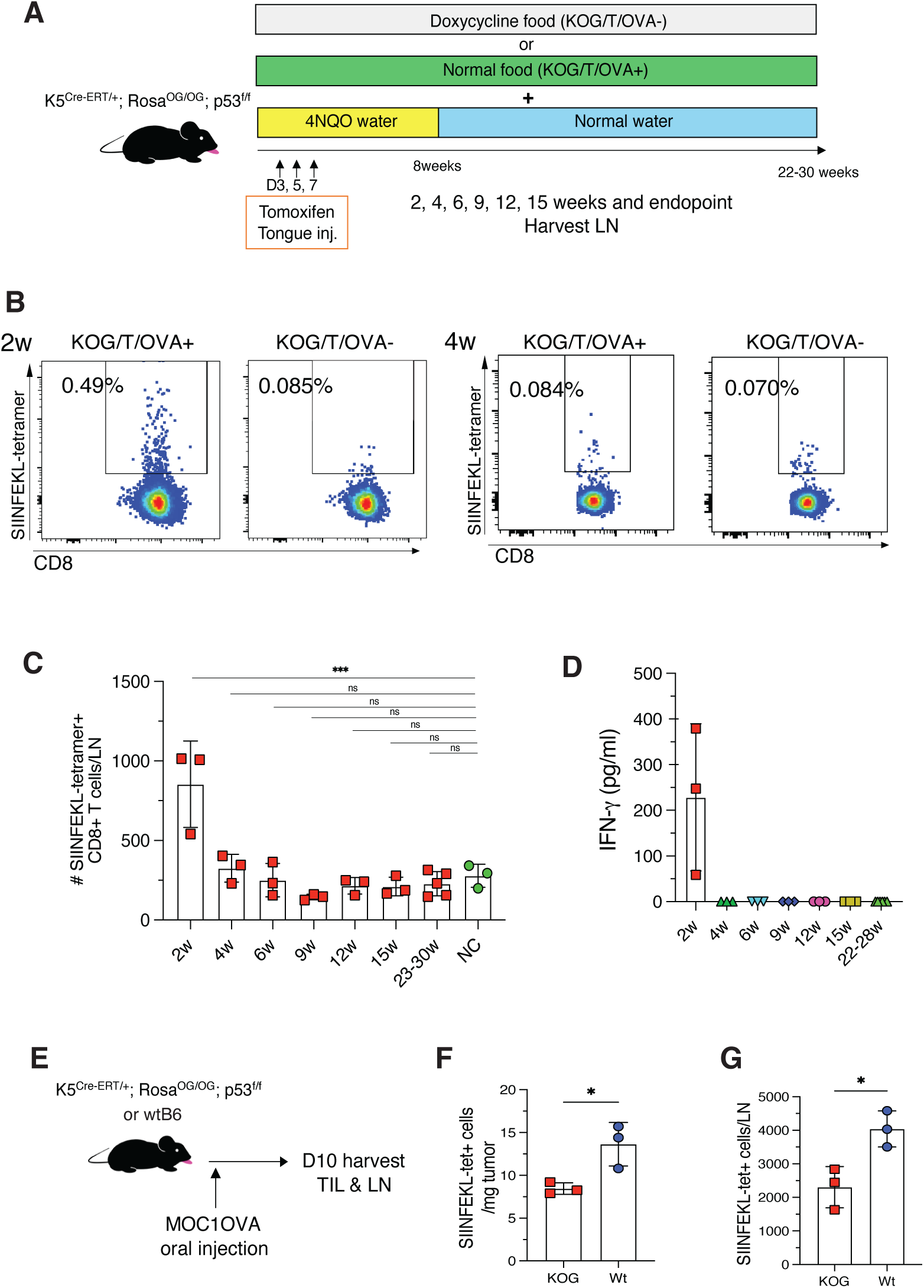
OVA-specific T cells were generated but subsequently became undetectable. A. Experimental schema for B-D. KOG mice were treated with 4NQO, doxycycline, and 4-OHT using the same protocol as in Figure 2A. Mice were sacrificed at specific time points or at the study endpoint. B. SIINFEKL-tetramer+ CD8+ T cells in DLNs from KOG/T/OVA^+^ mice 2 or 4 weeks after treatment initiation. Data were pre-gated on live cells, single cells, CD45+, CD3+, and CD8+ cells. LNs from doxycycline-treated mice were used as negative control of tetramer staining. Representative of two independent experiments (n=3). C. Quantification of the number of SIINFEKL-tetramer+ CD8+ T cells in DLNs from KOG/T/OVA^+^ mice over the time course. Mice were sacrificed at each time point and DLNs were analyzed. LNs from KOG/T/OVA^-^ were used as negative control of tetramer staining (NC). D. Quantification of IFN-γ production in supernatants of the cultures with lymphocytes from DLNs at each time points and SIINFEKL peptide (means ± SEM). DLNs were harvested at each time point and isolated lymphocytes were co-cultured with SIINFEKL peptide for 24 hours. The supernatants were assessed by IFN-γ ELISA. E. Experimental schema for F and G. 3 million MOC1OVA cells were inoculated into left buccal mucosa in KOG mice or wild type C57B/6 mice on day0. TILs and TDLNs were isolated on day10 and SIINFEKL-tetramer+ CD8+ T cells were analyzed by flow cytometry. F and G. SIINFEKL-tetramer+ CD8+ T cells per mg tumor in TILs (F) or per LN in TDLNs (G). *p<0.05, **p<0.01, ***p<0.001. Significance was evaluated by unpaired Student’s t-test and ANOVA with Tukey’s multiple comparison adjustment.

### Anti-PD-1 blockade did not restore OVA-specific T cell responses nor anti-tumor efficacy

We next examined the possible anti-tumor effects of anti-PD-1 blockade on 4NQO-induced KOG tumors. Anti-PD-1 antibody was injected twice weekly into tumor-bearing mice with or without OVA-expression (Figure 5A). Anti-PD-1 therapy did not result in tumor regression in either KOG/T/OVA^+^ or KOG/T/OVA^-^ tumors (Figure 5B-D). Moreover, CD8+ T cells and PD-1+ CD8+ T cells were not increased by anti-PD-1 blockade (Figure 5E and F). SIINFEKL-tetramer+ T cells were not detected in KOG/T/OVA^+^ tumors, even with anti-PD-1 blockade treatment. (Figure 5G). To examine whether OVA-specific T cells were initially present in an anergic state due to the PD-1 checkpoint, mice were treated with anti-PD-1 blockade at an earlier timepoint, when OVA-specific T cells were not detectable (Figure 5H). In this setting, anti-PD-1 inhibition also did not restore OVA-specific T cells (Figure 5I and J). These results suggested that KOG tumors were resistant to anti-PD-1 treatment regardless of OVA expression and anti-PD-1 blockade did not revive OVA-specific T cells.

**Figure 5:**
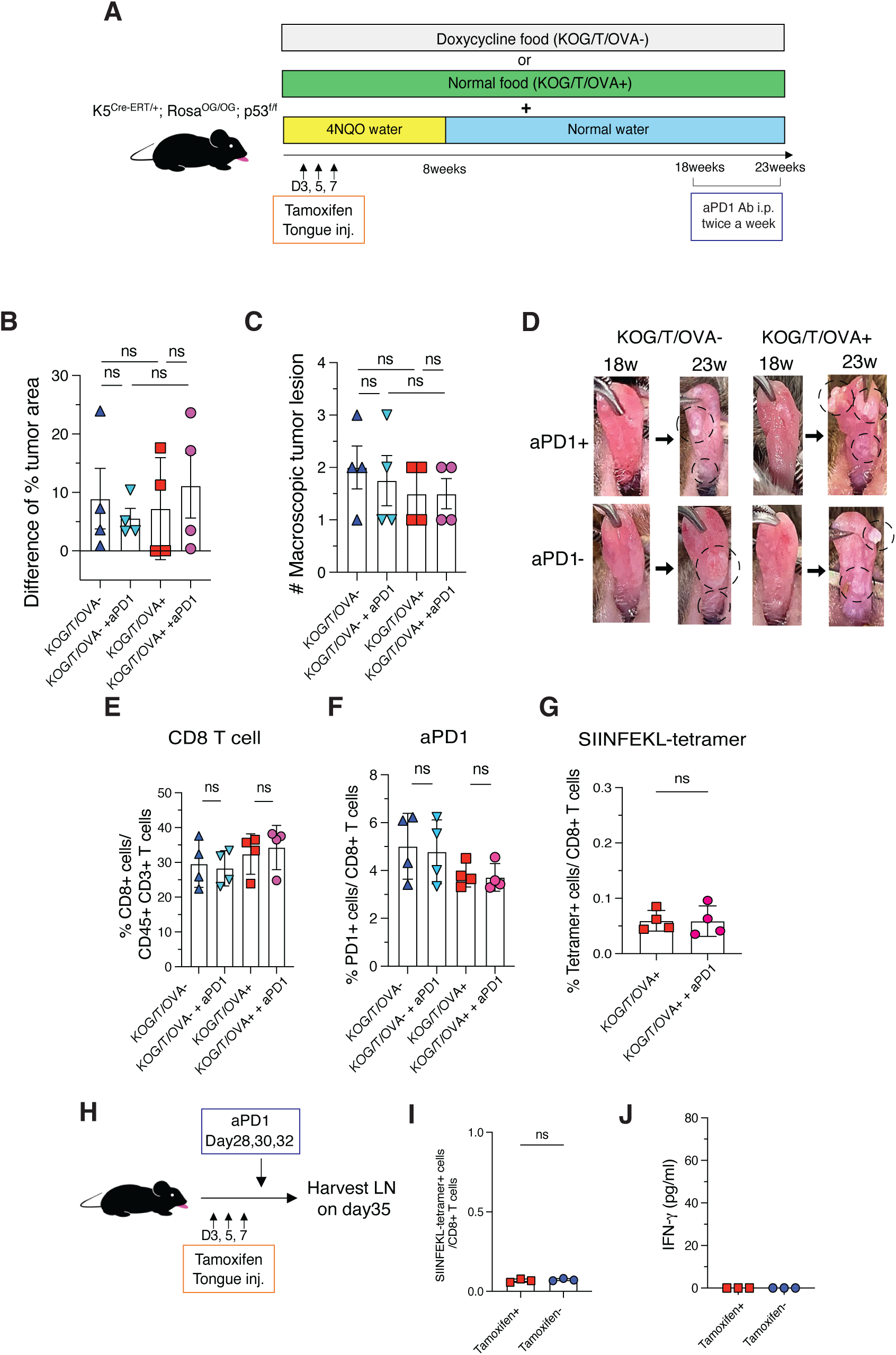
aPD-1 blockade did not restore OVA-specific T cells and show no anti-tumor effect. A. Experimental schema for the assessment of the anti-PD-1 anti-tumor effect. Treatment with 4NQO, doxycycline, and 4-OHT followed as in Figure 2A. anti-PD-1 Ab were injected twice a week starting at 18 weeks and tongues were harvested at 23 weeks. B. Difference in macroscopic tumor area before (18 weeks) and after (23 weeks) treatment (means ± SEM). C. The number of macroscopic tumor lesion at 23 weeks. D. Photographs of KOG/T/OVA^+^ and KOG/T/OVA^-^ tongues before (18 weeks) and after (23 weeks) anti-PD-1 therapy. E, F, G. The percentage of positive cells in TDLNs from 23 weeks tumor-bearing mice treated by aPD-1 blockade (means ± SEM). E. CD8+ T cells in CD45+ CD3+ T cells. F. PD-1+ cells in CD8+ T cells. G. SIINFEKL-tetramer+ cells in CD8+ T cells. H. Experimental schema for I and J. KOG mice were injected with 4-OHT on day3, 5, and 7 and anti-PD-1 on day28, 30, and 32. DLNs were harvested on day35. I. The percentage of SIINFEKL-tetramer+ cells in CD8+ T cells (means ± SEM). J. Quantification of IFN-γ production in supernatants of the cultures with lymphocytes from DLNs and SIINFEKL peptide (means ± SEM). *p<0.05, **p<0.01, ***p<0.001. Significance was evaluated by unpaired Student’s t-test and ANOVA with Tukey’s multiple comparison adjustment.

### Peptide vaccine restores OVA-specific T cells and attenuates tumor growth in combination with anti-PD-1

As impaired OVA specific T cell response was identified as a potential mechanism of immune evasion by OVA-expressing tumors in KOG mice, we next aimed to definitively examine if tumor control could be achieved with antigen specific T cells. Given the observed T cell response to MOC1-OVA, we tested whether OVA-specific T cells could be generated by OVA synthetic long peptide (SLP) and PolyI:C vaccine. Tumor-bearing mice were vaccinated with 3 doses of an OVA SLP in the buccal mucosa (Figure 6A). As shown in Figure 6B and C, this vaccine induced OVA-tetramer+ T cells in both KOG/T/OVA^+^ and KOG/T/OVA^-^ oral draining lymph nodes. OVA peptide co-culture and ELISA experiments confirmed that these T cells could recognize the OVA peptide effectively (Figure 6D).

**Figure 6:**
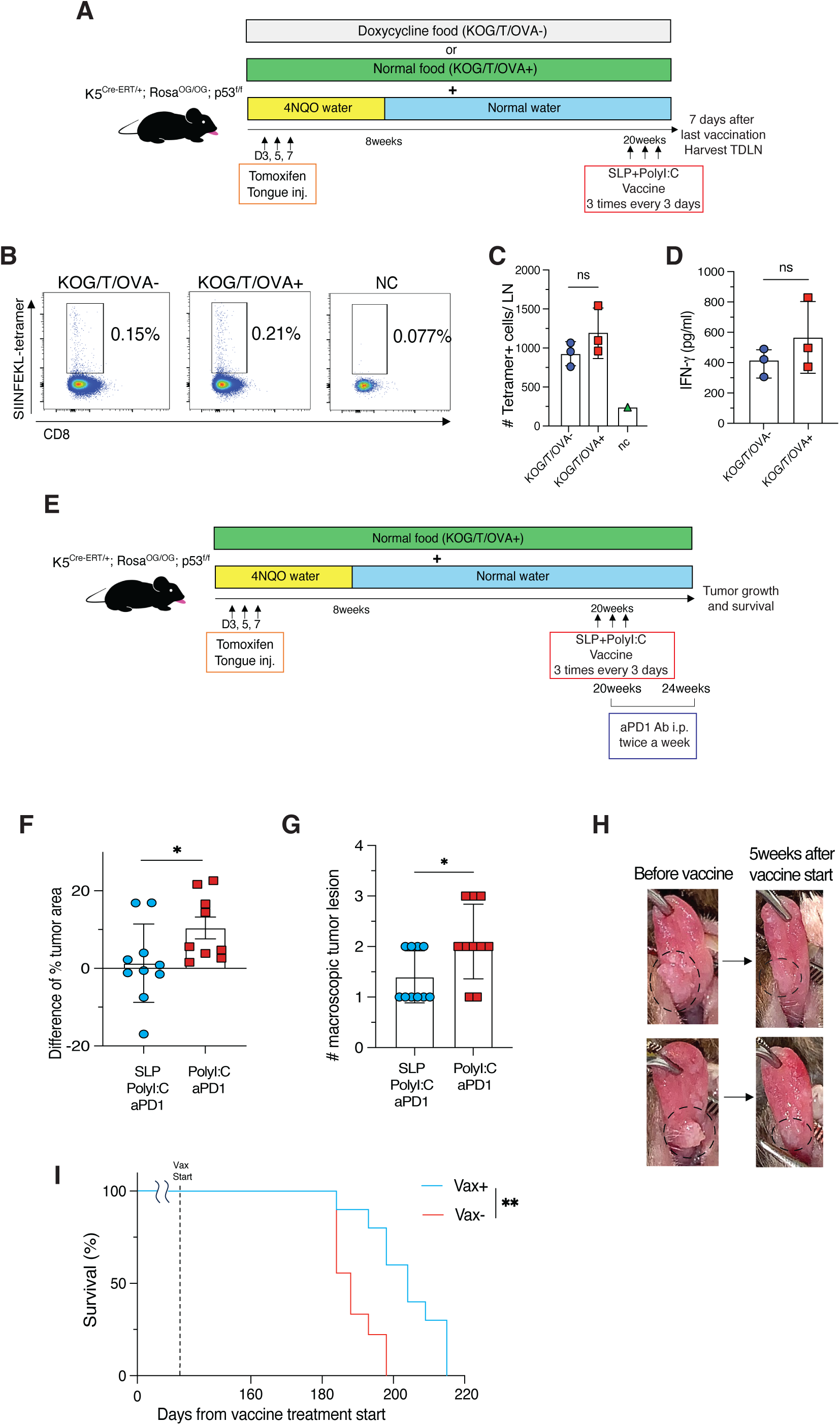
Peptide vaccine restores OVA-specific T cells and attenuates tumor growth in combination with anti-PD1. A. Experimental schema for B-D. Treatment of 4NQO, doxycycline, and 4-OHT followed the protocol in Figure 2A. 50µg OVA synthetic long peptide (SLP) containing SIINFEKL along with 100µg of polyI:C were injected into the buccal mucosa of tumor-bearing KOG mice three times every three days, starting at 20 weeks. TLDNs were harvested 7 days after last vaccine. B. SIINFEKL-tetramer+ CD8+ T cells in TDLNs from tumor-bearing KOG/T/OVA^+^ and KOG/T/OVA^-^ mice . LNs from untreated KOG mice were used as negative control of tetramer staining (NC). Representative of two independent experiments (n=3). C. Quantification of the number of SIINFEKL-tetramer+ CD8+ T cells in TDLNs (means ± SEM). D. Quantification of IFN-γ production in supernatants of the cultures with lymphocytes from TDLNs and SIINFEKL peptide (means ± SEM). Isolated lymphocytes from TDLNs were co-cultured with SIINFEKL peptide for 24 hours. Representative of two independent experiments. E. Experimental schema for F-I. Treatment of 4NQO, doxycycline, 4-OHT, and vaccine followed protocol outlined in Figure 6A. In addition, anti-PD-1 blockade was injected intraperitoneally twice a week and continued for 5 weeks. Treatment with PolyI:C injection plus aPD-1 therapy was used for control groups. F. Difference in macroscopic tumor area before (18 weeks) and after (23 weeks) treatment (means ± SEM).G. The number of macroscopic tumor lesion at 23 weeks. H. Photographs of the KOG/T/OVA^+^ mouse tongues treated with vaccine and anti-PD-1 blockade. I. Kaplan-Meier survival curve. *p<0.05, **p<0.01, ***p<0.001. Significance was evaluated by unpaired Student’s t-test and ANOVA with Tukey’s multiple comparison adjustment. Kaplan–Meier curve was used for survival.

Finally, we evaluated the anti-tumor efficacy of the peptide vaccine combined with anti-PD-1 therapy (Figure 6E). Remarkably, OVA SLP peptide and polyI:C vaccine with anti-PD-1 Ab significantly reduced the KOG tumor area compared to control group receiving PolyI:C plus anti-PD-1 therapy (Figure 5F-H). Moreover, overall survival was prolonged by OVA vaccine treatment (Figure 5I). Taken together, vaccination induced antigen-specific T cells in mice bearing oral tumor with immune evasion mechanisms involving impaired endogenous antigen-specific T cell response. This therapeutic approach resulted in tumor regression and extended survival.

## DISCUSSION

The carcinogen 4NQO triggers tumor genetic mutations similar to those found in human tobacco-related HNSCC (19) (20). Furthermore, 4NQO-induced malignancy also showed similar immune infiltration and response rate to anti-PD-1 therapy similar to those observed in human HNSCC (21). The advantage of the 4NQO model lies not only in its resemblance to human HNSCC, but also in its capability to observe tumorigenesis from its earliest stages. While the mutational burden is higher in invasive squamous cell carcinomas (SCCs) than hyperplasia and mild dysplasia, it is crucial to note that critical mutations in genes such as p53, Notch1, and Fat1 can already be detected in early-stage lesions (22). Recent studies of 4NQO-induced cancers at different pathological stages have also demonstrated that key immune-related genes increase from the dysplasia stage (23). In this study, we generated a 4NQO-induced HNSCC mouse model with genetically engineered epithelium-specific OVA expression. This unique model allows for the observation of immunoediting by endogenous antigen expression during tumorigenesis while maintaining HNSCC heterogeneity.

Other spontaneous cancers models have explored how antigen induction impacts tumor development, emphasizing its effect on eliciting tissue-specific immune responses. For example, in KPC mouse lung cancers with KRAS(G12D) activating mutation, p53 knockout and bearing the OG cassette, OVA expression led to tumor shrinkage (11). In contrast, in pancreatic cancer expression of this same antigen resulted in enhanced tumor growth due to cDC deficiency (11). Tissue-specific immune responses have been thoroughly investigated in the contexts of infectious diseases and autoimmunity, and their relevance to tumor immunity has also been established. These site specific responses also translate to ICI responses, as outcomes are markedly different depending on the site of metastasis (24). In preclinical syngeneic models, orthotopic transplantation has been shown to have a weaker immune surveillance relative to subcutaneous inoculation (25). However, in contrast to this finding, we observed that orthotopic oral inoculation elicits a stronger immune response and leads to tumor reduction compared to subcutaneous inoculation, again highlighting the difference in tissue-specific immune responses depending on cancer type and tumor site (26). To further understand HNSCC specific responses in the autochthonous setting, we engineered a HNSCC-specific preclinical model bearing the OG cassette. This approach aims to elucidate the role of the anti-tumor T cell immunity in HNSCC.

We observed that antigen-specific T cells were initially generated as a result of strong antigen expression from the early tumorigenesis but were consequently undetectable in peripheral tissues. With this absence of an antigen-specific response, we saw no difference in tumorigenesis compared to the antigen-free tumors and a similar absence of anti-PD-1 therapeutic efficacy. T cell absence is classified as central thymic tolerance during T cell development with subsequent peripheral loss (27). In this model, central tolerance to OVA-specific T cells may have developed presumably by leaky antigen expression in thymocytes. Leaky expression has been a known problem with the conventional inducible antigen engineered models because low level self-antigen expression can cause thymic tolerance (28, 29) (30). The recently described NINJA (inversion inducible joined neoantigen) model aims to prevent leaky neoantigen expression and tolerance (31, 32).

Since neoantigens are not expressed in normal tissues and development, the T cell repertoire maintains intact to mount anti-tumor responses to mutant allele products. Thus, our OG cassette-derived antigen most closely resembles tumor-associated antigens (TAAs) rather than neoantigens. Neoantigens have been extensively studied for their high immunogenicity but treatments utilizing personalized neoantigen-specific T cells are limited in their application across a broad patient population due to the unique tumor- and patient-specific characteristics. Thus, therapies targeting shared antigens, including TAAs, have been a focus of many studies (33) (34) (35). Notably, a detailed analysis of recent successful TIL therapy clinical trial in stage IV melanoma patients revealed that responding tumor-specific T cells targeted shared TAA-specific T cells and not neoantigen-specific T cells (36). This finding highlights the significance of researching TAA-specific T cells for advancing cancer treatment strategies.

In this study, central tolerance did not eliminate all OVA-specific T cells, however, upon antigen expression early in tumorigenesis, antigen-specific T cells were impaired in the periphery. Peripheral tolerance, as well as central tolerance, is crucial for preventing autoimmunity immune. Various mechanisms, both T cell intrinsic, such as anergy and clonal deletion, and T cell extrinsic, such as regulatory T cells, contribute to peripheral T cell tolerance (27, 37) (38). For instance, activated antigen-specific T cells in liver metastases were eliminated by FasL^+^ monocyte-derived macrophages, leading to acquired immunotherapy resistance (39). It is well recognized that checkpoint receptors, such as PD-1, play a role in promoting peripheral tolerance by causing the deletion or anergy of anti-tumor T cells (40, 41). Recent studies using the NINJA model revealed that PD-1 molecules result in peripheral tolerance, allowing the coexistence of antigen-expressing cells and antigen-specific T cells without an ongoing response (38). Our data that anti-PD-1 blockade did not restore antigen-specific T cells suggests that this mechanism was not occurring in our model system.

Despite the lack of an ongoing CD8+ T cell response, the therapeutic relevance of continued antigen expression in anti-PD-1 resistant tumors was highlighted by anti-tumor responses observed following vaccination against OVA. We found that that antigen-specific T cells, which were undetectable by tetramer and ex vivo peptide short stimulation, were induced by the vaccine, leading to tumor regression. These results suggest defining target antigens independent of endogenous antigen-specific T cell responses as a therapeutic strategy. Further investigation is needed to explore the specific TCRs recognizing the endogenous OG cassette expression versus those induced by vaccination. This study also supports the rationale for combining targeted immunotherapies, like cancer vaccines, with non-specific immunotherapies, such as anti-PD-1 inhibitors, for immune-cold tumors due to antigen-specific T cell deletion. This approach aligns with clinical evidence demonstrating the synergistic effects of combining cancer vaccines with anti-PD-1 inhibitors (42), suggesting a promising approach for enhancing therapeutic efficacy against these challenging tumor types including HNSCC. In conclusion, this study highlights a strategy for targeting tumors that evade the immune surveillance through impaired early T cell response in HNSCC.

## DECLARATIONS

### Ethics approval

All experimental animal procedures were approved by the institutional animal care and use committee (IACUC) of the Dana-Farber Cancer Institute.

### Author contributions

Conceptualization, M.K. and R.U.; Methodology, M.K., A.M.E., and R.U.; Investigation, M.K., M.R., J.Q., S.S., H.K., and S.B.W.; Writing – Original Draft, M.K. and R.U.; Writing –Review & Editing, All authors; Funding Acquisition, R.U.; Resources, A.M.E. and R.U.; Supervision, A.M.E., and R.U.

## Acknowledgements

We thank Dr. David Denardo for providing the OG mouse and Dr. Clint Allen for supplying reagents. We thank the NIH tetramer core for provision of reagents and all members of the Uppaluri lab for discussions.

## REFERENCES

1. Ruffin AT, Li H, Vujanovic L, Zandberg DP, Ferris RL, and Bruno TC. Improving head and neck cancer therapies by immunomodulation of the tumour microenvironment. Nat Rev Cancer. 2023;23(3):173–88.

2. Gillison ML, Blumenschein G, Fayette J, Guigay J, Colevas AD, Licitra L, et al. Long-term Outcomes with Nivolumab as First-line Treatment in Recurrent or Metastatic Head and Neck Cancer: Subgroup Analysis of CheckMate 141. Oncologist. 2022;27(2):e194–e8.

3. Schreiber RD, Old LJ, and Smyth MJ. Cancer immunoediting: integrating immunity’s roles in cancer suppression and promotion. Science. 2011;331(6024):1565-70.

4. Hayman TJ, Baro M, MacNeil T, Phoomak C, Aung TN, Cui W, et al. STING enhances cell death through regulation of reactive oxygen species and DNA damage. Nature communications. 2021;12(1):2327.

5. Comprehensive genomic characterization of head and neck squamous cell carcinomas. Nature. 2015;517(7536):576–82.

6. Grandis JR, Falkner DM, Melhem MF, Gooding WE, Drenning SD, and Morel PA. Human leukocyte antigen class I allelic and haplotype loss in squamous cell carcinoma of the head and neck: clinical and immunogenetic consequences. Clinical cancer research : an official journal of the American Association for Cancer Research. 2000;6(7):2794–802.

7. López-Albaitero A, Nayak JV, Ogino T, Machandia A, Gooding W, DeLeo AB, et al. Role of antigen-processing machinery in the in vitro resistance of squamous cell carcinoma of the head and neck cells to recognition by CTL. Journal of immunology. 2006;176(6):3402–9.

8. Clavijo PE, Moore EC, Chen J, Davis RJ, Friedman J, Kim Y, et al. Resistance to CTLA-4 checkpoint inhibition reversed through selective elimination of granulocytic myeloid cells. Oncotarget. 2017;8(34):55804–20.

9. Sun L, Clavijo PE, Robbins Y, Patel P, Friedman J, Greene S, et al. Inhibiting myeloid-derived suppressor cell trafficking enhances T cell immunotherapy. JCI Insight. 2019;4(7).

10. Saito S, Kono M, Nguyen HCB, Egloff AM, Messier C, Lizotte P, et al. Targeting Dendritic Cell Dysfunction to Circumvent anti-PD1 Resistance in Head and Neck Cancer. Clin Cancer Res. 2024.

11. Hegde S, Krisnawan VE, Herzog BH, Zuo C, Breden MA, Knolhoff BL, et al. Dendritic Cell Paucity Leads to Dysfunctional Immune Surveillance in Pancreatic Cancer. Cancer Cell. 2020;37(3):289–307.e9.

12. Wu JS, Zheng M, Zhang M, Pang X, Li L, Wang SS, et al. Promotes 4-Nitroquinoline-1-Oxide-Induced Oral Carcinogenesis With an Alteration of Fatty Acid Metabolism. Front Microbiol. 2018;9:2081.

13. Carper MB, Troutman S, Wagner BL, Byrd KM, Selitsky SR, Parag-Sharma K, et al. An Immunocompetent Mouse Model of HPV16(+) Head and Neck Squamous Cell Carcinoma. Cell Rep. 2019;29(6):1660–74.e7.

14. Karakasheva TA, Kijima T, Shimonosono M, Maekawa H, Sahu V, Gabre JT, et al. Generation and Characterization of Patient-Derived Head and Neck, Oral, and Esophageal Cancer Organoids. Curr Protoc Stem Cell Biol. 2020;53(1):e109.

15. Judd NP, Winkler AE, Murillo-Sauca O, Brotman JJ, Law JH, Lewis JS, Jr., et al. ERK1/2 Regulation of CD44 Modulates Oral Cancer Aggressiveness. Cancer research. 2012;72(1):365–74.

16. Onken MD, Winkler AE, Kanchi KL, Chalivendra V, Law JH, Rickert CG, et al. A surprising cross-species conservation in the genomic landscape of mouse and human oral cancer identifies a transcriptional signature predicting metastatic disease. Clinical cancer research : an official journal of the American Association for Cancer Research. 2014;20(11):2873–84.

17. Kono M, Saito S, Egloff AM, Allen CT, and Uppaluri R. The mouse oral carcinoma (MOC) model: A 10-year retrospective on model development and head and neck cancer investigations. Oral oncology. 2022;132:106012.

18. Sun L, Moore E, Berman R, Clavijo PE, Saleh A, Chen Z, et al. WEE1 kinase inhibition reverses G2/M cell cycle checkpoint activation to sensitize cancer cells to immunotherapy. Oncoimmunology. 2018;7(10):e1488359.

19. Bhatia S, Oweida A, Lennon S, Darragh LB, Milner D, Phan AV, et al. Inhibition of EphB4-Ephrin-B2 Signaling Reprograms the Tumor Immune Microenvironment in Head and Neck Cancers. Cancer Res. 2019;79(10):2722–35.

20. Cao Y, Dong H, Li G, Wei H, Xie C, Tuo Y, et al. Temporal and spatial characteristics of tumor evolution in a mouse model of oral squamous cell carcinoma. BMC Cancer. 2022;22(1):1209.

21. Wang Z, Wu VH, Allevato MM, Gilardi M, He Y, Luis Callejas-Valera J, et al. Syngeneic animal models of tobacco-associated oral cancer reveal the activity of in situ anti-CTLA-4. Nature communications. 2019;10(1):5546.

22. Sequeira I, Rashid M, Tomás IM, Williams MJ, Graham TA, Adams DJ, et al. Genomic landscape and clonal architecture of mouse oral squamous cell carcinomas dictate tumour ecology. Nat Commun. 2020;11(1):5671.

23. Lee YM, Hsu CL, Chen YH, Ou DL, Hsu C, and Tan CT. Genomic and Transcriptomic Landscape of an Oral Squamous Cell Carcinoma Mouse Model for Immunotherapy. Cancer Immunol Res. 2023;11(11):1553–67.

24. Tumeh PC, Hellmann MD, Hamid O, Tsai KK, Loo KL, Gubens MA, et al. Liver Metastasis and Treatment Outcome with Anti-PD-1 Monoclonal Antibody in Patients with Melanoma and NSCLC. Cancer Immunol Res. 2017;5(5):417–24.

25. Horton BL, Morgan DM, Momin N, Zagorulya M, Torres-Mejia E, Bhandarkar V, et al. Lack of CD8(+) T cell effector differentiation during priming mediates checkpoint blockade resistance in non-small cell lung cancer. Sci Immunol. 2021;6(64):eabi8800.

26. Kono M, Saito S, Rokugo M, Egloff AM, and Uppaluri R. Enhanced oral versus flank lymph node T cell response parallels anti-PD1 efficacy in head and neck cancer. Oral Oncol. 2024;152:106795.

27. Ghorani E, Swanton C, and Quezada SA. Cancer cell-intrinsic mechanisms driving acquired immune tolerance. Immunity. 2023;56(10):2270–95.

28. DuPage M, Mazumdar C, Schmidt LM, Cheung AF, and Jacks T. Expression of tumour-specific antigens underlies cancer immunoediting. Nature. 2012;482(7385):405-9.

29. Probst HC, Lagnel J, Kollias G, and van den Broek M. Inducible transgenic mice reveal resting dendritic cells as potent inducers of CD8+ T cell tolerance. Immunity. 2003;18(5):713–20.

30. Malhotra D, Linehan JL, Dileepan T, Lee YJ, Purtha WE, Lu JV, et al. Tolerance is established in polyclonal CD4(+) T cells by distinct mechanisms, according to self-peptide expression patterns. Nat Immunol. 2016;17(2):187–95.

31. Fitzgerald B, Connolly KA, Cui C, Fagerberg E, Mariuzza DL, Hornick NI, et al. A mouse model for the study of anti-tumor T cell responses in Kras-driven lung adenocarcinoma. Cell Rep Methods. 2021;1(5).

32. Damo M, Fitzgerald B, Lu Y, Nader M, William I, Cheung JF, et al. Inducible de novo expression of neoantigens in tumor cells and mice. Nat Biotechnol. 2021;39(1):64–73.

33. Dreno B, Thompson JF, Smithers BM, Santinami M, Jouary T, Gutzmer R, et al. MAGE-A3 immunotherapeutic as adjuvant therapy for patients with resected, MAGE-A3-positive, stage III melanoma (DERMA): a double-blind, randomised, placebo-controlled, phase 3 trial. Lancet Oncol. 2018;19(7):916–29.

34. Kjeldsen JW, Lorentzen CL, Martinenaite E, Ellebaek E, Donia M, Holmstroem RB, et al. A phase 1/2 trial of an immune-modulatory vaccine against IDO/PD-L1 in combination with nivolumab in metastatic melanoma. Nat Med. 2021;27(12):2212–23.

35. Pant S, Wainberg ZA, Weekes CD, Furqan M, Kasi PM, Devoe CE, et al. Lymph-node-targeted, mKRAS-specific amphiphile vaccine in pancreatic and colorectal cancer: the phase 1 AMPLIFY-201 trial. Nat Med. 2024;30(2):531–42.

36. Dolton G, Rius C, Wall A, Szomolay B, Bianchi V, Galloway SAE, et al. Targeting of multiple tumor-associated antigens by individual T cell receptors during successful cancer immunotherapy. Cell. 2023;186(16):3333–49.e27.

37. ElTanbouly MA, and Noelle RJ. Rethinking peripheral T cell tolerance: checkpoints across a T cell’s journey. Nat Rev Immunol. 2021;21(4):257–67.

38. Damo M, Hornick NI, Venkat A, William I, Clulo K, Venkatesan S, et al. PD-1 maintains CD8 T cell tolerance towards cutaneous neoantigens. Nature. 2023;619(7968):151-9.

39. Yu J, Green MD, Li S, Sun Y, Journey SN, Choi JE, et al. Liver metastasis restrains immunotherapy efficacy via macrophage-mediated T cell elimination. Nat Med. 2021;27(1):152–64.

40. Nishimura H, Nose M, Hiai H, Minato N, and Honjo T. Development of lupus-like autoimmune diseases by disruption of the PD-1 gene encoding an ITIM motif-carrying immunoreceptor. Immunity. 1999;11(2):141–51.

41. Probst HC, McCoy K, Okazaki T, Honjo T, and van den Broek M. Resting dendritic cells induce peripheral CD8+ T cell tolerance through PD-1 and CTLA-4. Nat Immunol. 2005;6(3):280–6.

42. Ott PA, Hu-Lieskovan S, Chmielowski B, Govindan R, Naing A, Bhardwaj N, et al. A Phase Ib Trial of Personalized Neoantigen Therapy Plus Anti-PD-1 in Patients with Advanced Melanoma, Non-small Cell Lung Cancer, or Bladder Cancer. Cell. 2020;183(2):347–62 e24.

