## Supplemental for "Cancer vaccine attenuates carcinogen induced head and neck cancer with impaired early T cell response"

**Supplemental Figure**

S1. Representative of the measurement of tumor area per tongue.

S2A. Gating strategy of GFP+ CD45- tongue epithelium cells.

S2B. Gating strategy of CFSE+ CD8 + T cells in LNs.

S3A. Gating strategy of SIINFEKL-tetramer+ CD8+ T cells in LNs.

S3B. OVA-tetramer+ CD4+ T cells in 2-week and 4-week KOG/T/$OVA^{+}$ mice. Data were pre-gated on live cells, single cells, CD45+, CD3+ and CD4+ cells. Representative of three independent experiments. Left is representative data and right is quantification (means± SEM)

S3C. CD40 agonist was injected to KOG/T/$OVA^{+}$ mice intraperitoneally on day14 and SIINFEKL-tetramer+ CD8+ T cells were evaluated by flow cytometry. Data were pre-gated on live cells, single cells, CD45+, CD3+ and C84+ cells. Representative of four mice.

Table S1. Antibody lists

FigureS1

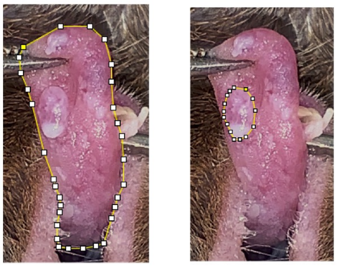

Whole tongue    Tumor area

FigureS2

A

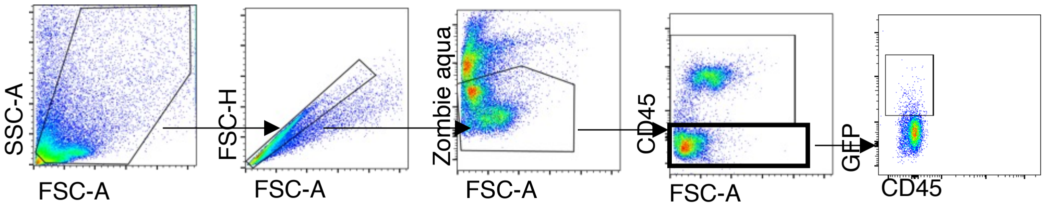

SSC-A
FSC-A
FSC-H
FSC-A
Zombie aqua
FSC-A
CD45
FSC-A
GFP
CD45


B

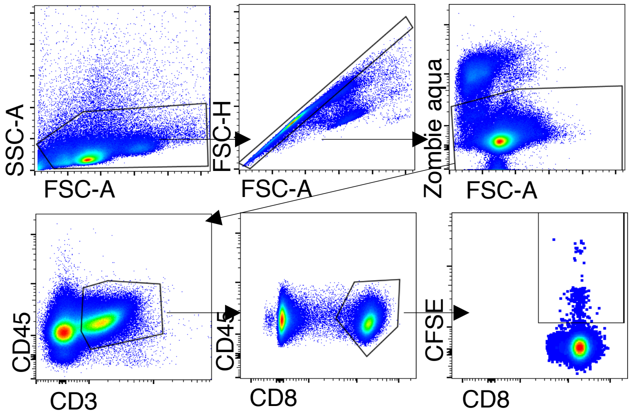

SSC-A
FSC-A
FSC-H
FSC-A
Zombie aqua
FSC-A
CD45
CD3
CD45
CD8
CFSE
CD8

FigureS3

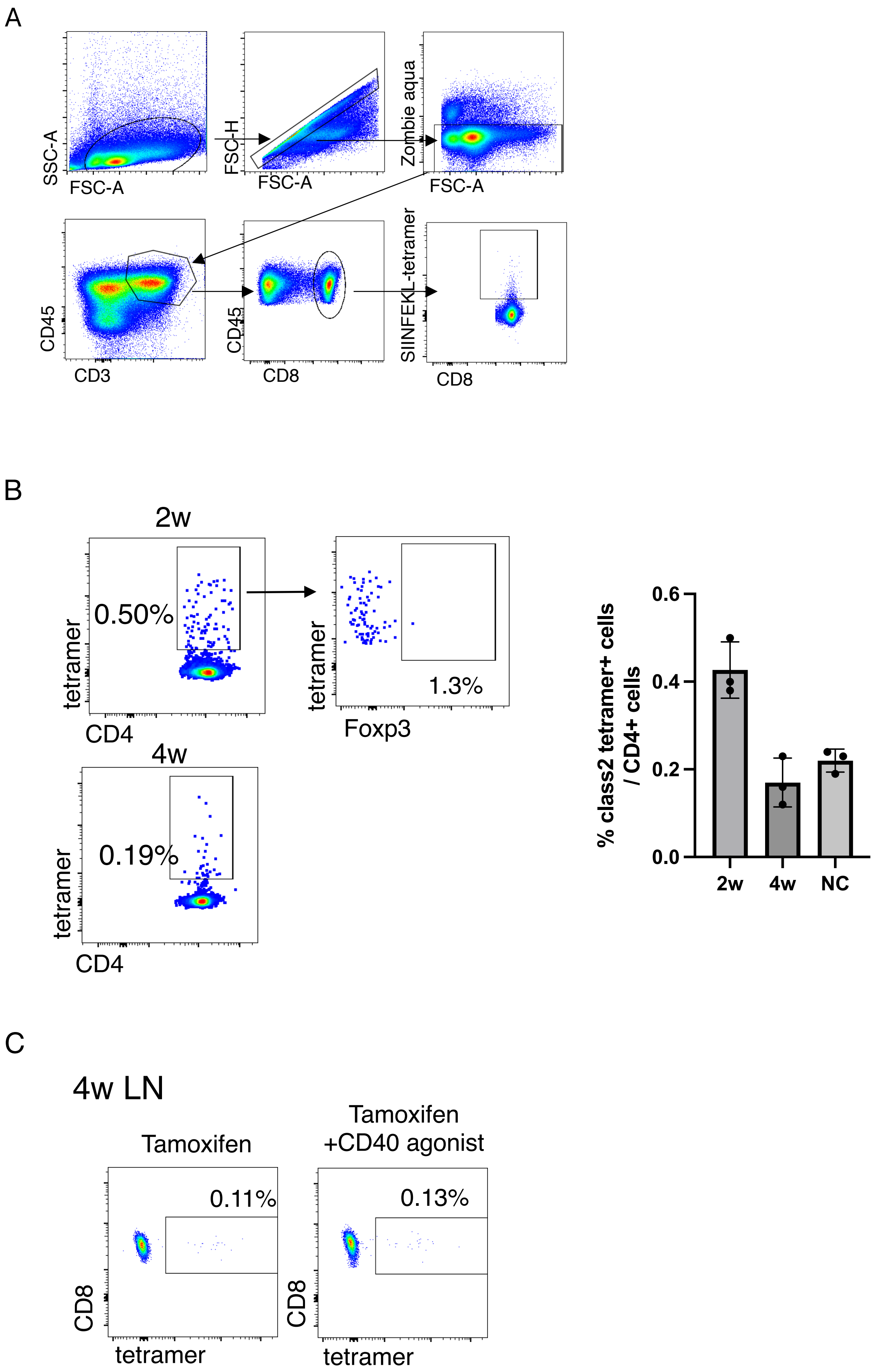
A
SSC-A
FSC-A
FSC-H
FSC-A
Zombie aqua
FSC-A
CD45
CD3
CD45
CD8
SIINFEKL-tetramer
CD8
B
2w
0.50%
tetramer
CD4
tetramer
1.3%
Foxp3
4w
0.19%
tetramer
CD4
% class2 tetramer+ cells / CD4+ cells
0.6
0.4
0.2
0.0
2w
4w
NC
C
4w LN
Tamoxifen
Tamoxifen +CD40 agonist
0.11%
0.13%
CD8
tetramer
CD8
tetramer

**TableS1**

| | | |
|---|---|---|
| Anti-mouse CD3 APC/Cyanine7 (clone 145-2C11) | BioLegend | Cat#100330 |
| Anti-mouse CD3 BV650 (clone 17A2) | BioLegend | Cat#100229 |
| Anti-mouse CD3 PE (clone 145-2C11) | BioLegend | Cat#100308 |
| Anti-mouse CD4 Alexa Fluor700 (clone RM4-4) | BioLegend | Cat#116021 |
| Anti-mouse CD4 FITC (clone GK1.5) | BioLegend | Cat#100405 |
| Anti-mouse CD45.2 BV421 (clone 104) | BioLegend | Cat#109832 |
| Anti-mouse CD45.2 PerCp (clone 104) | BioLegend | Cat#109825 |
| Anti-mouse CD45.2 BV711 (clone 104) | BioLegend | Cat#109847 |
| Anti-mouse CD8 BV421 (clone 53-6.7) | BioLegend | Cat#100737 |
| Anti-mouse CD8 BV605 (clone 53-6.7) | BioLegend | Cat#100743 |
| Anti-mouse CD8 FITC (clone KT15) | MBL | Code No.K0227-4 |
| Anti-mouse PD1 APC (clone 29F.1A12) | BioLegend | Cat#135210 |
| Anti-mouse H-2Kb bound to SIINFEKL (clone 25-D1.16 ) | BioLegend | Cat#141603 |
